# Development of Nanobodies as first-in-class inhibitors for the NEDP1 deNEDDylating enzyme

**DOI:** 10.1101/2020.03.20.999326

**Authors:** Naima Abidi, Helene Trauchessec, Gholamreza Hassanzadeh-Ghassabeh, Martine Pugniere, Serge Muyldermans, Dimitris P. Xirodimas

## Abstract

Protein NEDDylation emerges as an important post-translational modification and an attractive target for therapeutic intervention. Modification of NEDD8 onto substrates is finely balanced by the co-ordinated activity of conjugating and deconjugating enzymes. The NEDP1/DEN1/SENP8 protease is a NEDD8 specific processing and deconjugating enzyme that regulates the NEDDylation mainly of non-cullin substrates. Here, we report the development and characterisation of nanobodies as first-in-class inhibitors for NEDP1. The nanobodies display high-affinity (low nM) against NEDP1 and specifically inhibit NEDP1 processing activity *in vitro* and NEDP1 deconjugating activity in tissue-culture cells and in cell extracts. We also isolated nanobodies that bind to NEDP1 with high-affinity but do not affect NEDP1 activity. The developed nanobodies provide new tools to study the function of NEDP1 and to prevent deNEDDylation in cell extracts used in biochemical assays.

## Introduction

Modification of proteins with the ubiquitin-like molecule NEDD8 is essential almost in all tested organisms. Similarly to the modification with ubiquitin, protein NEDDylation depends on the activity of the so-called E1, E2 and E3 enzymes^1,2^. Prior to its activation, NEDD8 has to be processed to expose the C-terminal diGlycine motif. While several enzymes with dual activity towards ubiquitin and NEDD8 have been reported as NEDD8 processing enzymes, the NEDP1 (DEN1, SENP8) cysteine protease has specific NEDD8 processing activity^1,2^. Upon its processing, NEDD8 is activated by a single E1 enzyme (heterodimer Uba3/APPBP1) and conjugated through the activity of two E2 conjugating enzymes (Ube2M, Ube2F) and multiple E3-ligases^1,2^. The activation and conjugation of NEDD8 by the above-described specific NEDD8 enzymes defines the so-called canonical pathway. However, under conditions of proteotoxic stress, in addition to the conjugation of NEDD8 by the canonical pathway, NEDD8 is activated and conjugated by enzymes of the ubiquitin system, leading to the formation of hybrid ubiquitin-NEDD8 chains^3,4^. This defines the so-called atypical pathway that is involved in the formation of nuclear protein aggregates and control of the nuclear Ubiquitin Proteasome System^5,6^.

The action of NEDD8 conjugating enzymes is finely balanced by the activity of deconjugating (deNEDDylating) enzymes. Similarly to NEDD8 processing, several proteases with dual activity towards ubiquitin and NEDD8 can promote NEDD8 deconjugation from substrates^1,2^. However, the two specific deNEDDylating enzymes are the COP9 signalosome and NEDP1. COP9, a zinc metalloprotease, has minimal affinity towards NEDD8 but specifically deconjugates NEDD8 from cullins, the best established target for NEDD8^1,2^. NEDP1 on the other hand displays deNEDDylating activity towards non-cullin substrates^7–18^ and its deletion causes a dramatic increase in global protein NEDDylation^11,12,19,20^, with the accumulation of poly-NEDD8 chains through lysines K11/K48 as a prominent effect^11,19^. Importantly, both deNEDDylating activities control the canonical NEDD8 pathway^1,2,19^. While the function of the COP9 signalosome as a key component of the NEDD8 pathway through regulation of the Cullin Ring Ligases has been established, the role of NEDP1 in homeostatic and stress responses is only now emerging^1,2^. Recent studies indicated a role for NEDP1 in tumour progression; The upregulation of NEDD8 in hepatocellular carcinoma was related to a decrease in the levels of NEDP1. Importantly, decrease in NEDP1 levels provides resistance of tumour cells to DNA damage induced apoptosis^9, 19–21^.

Targeting the NEDD8 pathway is a promising therapeutic approach. Inhibitors for the NEDD8 E1 enzyme (MLN4924/Pevonedistat) are in Phase III clinical trials for cancer treatment^1,22,23^. The promising anti-tumour effects of MLN4924 have generated an interest in the field and additional components of the NEDD8 pathway including E2 conjugating enzymes, E3-ligases and the COP9 signalosome have been targeted^1,24–26^. Apart for their potential clinical testing and therapeutic use, such compounds provide invaluable tools to study the function of the target in question. However, specific inhibitors of the deNEDDylating enzyme NEDP1 do not currently exist.

Nanobodies (Nbs) are the smallest antigen binding fragments, with molecular mass of 15kDa, containing the full antigen-binding capacity of the original heavy chain-only antibody, unique for camelids. In addition to the benefits of conventional antibodies, such as high affinity and selectivity for a target, easy tailoring into pluripotent constructs and low inherent toxicity, Nbs have technological and biophysical advantages that outperform conventional antibodies; 1. Nbs are only a tenth of the size of a conventional antibody able to penetrate tissues more effectively than conventional antibodies and they can recognize uncommon or hidden epitopes^27,28^ 2. They are naturally soluble in aqueous solution and do not have a tendency to aggregate^29^. 3. They can be easily expressed in cells as fusion proteins enabling the recognition of the target of interest in living cells. These features have already lead to a number of biotechnological and medical applications in which Nbs excel other antibody formats^30,31^.

Here, we describe the development and characterisation of Nbs that specifically inhibit NEDP1 activity. Using *in vitro* and *in vivo* assays we show that the developed Nbs have a high affinity (low nM scale) for human NEDP1 and potently inhibit both the processing and deconjugating activity of NEDP1. Expression of the Nbs in human cells in tissue culture cause the accumulation of NEDD8 conjugates equivalent to the level observed upon NEDP1 deletion by CRISPR/Cas9. In summary, the study describes the development of Nbs as first-in-class inhibitors for NEDP1 that should provide invaluable tools to study the function of a key deNEDDylating activity.

## Materials and Methods

### Reagents

All common chemicals were purchased from Sigma Aldrich. Protease Inhibitor Cocktail Tablets EDTA-free (Roche). Rabbit monoclonal anti-NEDD8 (1:2000), Y297 (GeneTex, GTX61205), rabbit anti-ubiquitin (1:2000), (DAKO, Z0458), mouse anti-GAPDH (1:5000) (6C5, ab8245), mouse anti-SUMO-2 (1:2000) (gift from Dr Guillaume Bossis), sheep anti-NEDP1 (1:1000) (gift from Prof. Ronald Hay), mouse anti-Flag (1:2000), (Sigma, F1804), mouse anti-His6 (Clontech, 631212). Anti-Flag beads were purchased from SIGMA.

### Cell culture

U2OS cells were grown in Dulbecco’s modified Eagle’s medium (DMEM) supplemented with 10% fetal bovine serum (FBS) and antibiotics (Penicillin, 50U/ml and Streptomycin 50μg/ml) in 5% CO2 at 37°C in a humidified incubator. Cell lines were routinely tested for mycoplasma contamination but have not been authenticated. Cells were kept in culture for a maximum of 20 passages. Cells stably expressing Nbs were selected with G418 (1mg/ml).

### Immunisation of llama, construction of anti-NEDP1 Nb library and selection of specific antibody fragments (Biopanning)

A llama was injected subcutaneously by 140μg recombinant human NEDP1 protein emulsified with Gerbu adjuvant, six times at weekly intervals. After the last immunisation, 50ml of blood were taken and peripheral blood lymphocytes (PBLs) were extracted with lympho-prep (Nycomed, Switzerland). Library construction was done as previously described^32,33^. Briefly, Trizol-extracted mRNA from the pellet was used as a template for RT-PCR to generate first strand cDNA using oligo-dT primers. cDNA was used as template to amplify 2 different products using specific primers for the variable domain until the CH2 domain CALL001 (5’-GTCCTGGCTCTCTTCTACAAGG) and CALL002 (5’-GGTACGTGCTGTTGAACTGTTCC) to amplify the heavy chain antibody gene fragments from the variable region to the CH2 region, which yielded the VH and the VHH containing gene fragments of 900 and 600bp, respectively. The 600bp product was excised from 1% agarose gel, used as template for a second PCR via nested primers VHH-For primer 38 (5’-GGA CTA GTG CGG CCG CTG GAG ACG GTG ACC TGG GT) and VHH-Back primer A6E (5’-GAT GTG CAG CTG CAG GAG TCT GGR GGA GG)^32^ to amplify the VHH sequence of about 400bp and digested with Not I and PstI restriction enzymes (NEB). The PCR fragments were ligated into the phagemid vector pHEN4^34^ and transformed into electro-competent *E. coli* TG1 cells. The resulting Nb library was infected with M13K07 helper phages for the expression of Nbs on the phages. Biopanning was performed by coating ELISA plates (Nunc) with 20μg of recombinant NEDP1 per/well. Phage enrichment was achieved by 3 rounds of *in vitro* selection. Phages were eluted as described^33^. TG1 *E.coli* cells were infected with the eluted phages, and selected from LB ampicillin plates. One hundred viable colonies were randomly picked from each round of panning and their VHH was expressed as described^33^. The periplasmic extracts (PE) were obtained through osmotic shock as described^35^. The enrichment of each round of panning was checked by a polyclonal phage ELISA with anti-M13-HRP antibodies.

### Expression and purification of antibody fragments for ELISA testing

The cloned insert expressing Nbs recognizing NEDP1 recombinant protein, was sequenced using the RP or GIII primer and sequences were grouped based on the differences in their complementarity determining regions (CDRs). Representatives of each group were recloned into the expression vector pHEN6 using BstEII and PstI (NEB) enzymes. The plasmid constructs were transformed into WK6 *E. coli* cells and large quantities of His6 tagged recombinant Nbs were purified with 1ml of HIS-Select Nickel Affinity Gel (Sigma-Aldrich) following periplasmic expression through osmotic shock. Elution was performed with 1 column volume of 0.5M imidazole, repeated 3 times. The Nbs yields after purification varied from 1 to 15mg/L of *E. coli* culture. Solid phase ELISA was performed by coating recombinant NEDP1 (20μg/well) on Maxisorb 96 well plates (Nunc) at 4°C overnight. The ELISA was performed using purified Nbs as primary antibodies, followed by an anti-His_6_ and an anti-mouse antibody conjugated to horseradish peroxidase (Sigma-Aldrich). A non-relevant Nb (NbAahI)^36^ was used as negative control for antibody detection while a non-relevant protein, Bovine Serum Albumin (BSA) was used as negative control for NEDP1 coating.

### Cross-reactivity assay

The specificity of the 17 purified Nbs was tested by indirect ELISA, and 3 different proteins, NEDP1, ULP3 (the C. elegans homologue of NEDP1) and GST were chosen to test potential cross-reactivity. The same amount of the 3 proteins (20μg) was used to coat a 96-well plate, incubated overnight at 4°C. After blocking, purified Nbs were incubated with antigen at 37°C for 1h. Then, anti-His_6_ (diluted 1:2000) was added and incubated at 37°C for 1h. After another washing step, an anti-mouse antibody conjugated to horseradish peroxidase (Sigma-Aldrich) was added to the plate and incubated at 37°C for 1h. The reaction was stopped by the addition of 50μl of 3M H_2_SO_4_, and then the absorbance at 492nm was read on a microtiter plate reader.

The specificity of anti-NEDP1 Nbs was evaluated by monitoring their interaction against recombinant human NEDP1, ULP-3 and GST as GST-NEDP1 was used for immunisation. Only candidates with no cross reactivity to ULP-3 and GST, were selected to measure their affinity by surface plasmon resonance.

### Surface Plasmon Resonance (SPR) analysis

Nbs affinities for NEDP1 were determined on a Biacore 2000 apparatus (Biacore AB, Uppsala, Sweden). Nanobodies (355nM) were immobilized on anti-His_6_ antibody covalently immobilized on a carboxymethyl dextran sensorchip (CM5 Biacore AB) using the amine coupling method according to the manufacturer’s instructions. The immobilization level was 500 resonance units (RU). Different concentrations of NEDP1 ranging from 0.75 to 200nM in HEPES-buffered saline (0.01M HEPES, pH 7.4, 0.15M NaCl, 3mM EDTA, 0.005% surfactant P20) were injected and exposed to the Nb. All sensorgrams were fitted by subtracting the signal from the reference flow cell and were treated using BIAevaluation version 3.2 software (Biacore AB).

### Expression and Purification of Recombinant Proteins

His_6_-MBP-NEDD8-Ub and GST-NEDP1 (wild type and catalytic C163A mutant) were expressed and purified from, *E coli* BL21 (DE3) as previously described^37,38^. Expression of the protein was confirmed by western blotting, and the activity of recombinant protein was tested by *in vitro* deconjugation and processing assays.

### *in vitro* deconjugation assays

To determine the deconjugation activity of NEDP1, an increasing amount of the recombinant enzyme was incubated with 20μg of extracts from U2OS NEDP1 knockout cells (H6) that have high levels of NEDD8 conjugates^19^ at 37°C for 30min in 15μl of reaction buffer containing 50mM Tris–HCl (pH 8.0), 50mM NaCl and 5mM b-mercaptoethanol. The reaction was stopped by the addition of 3μl of 6X SDS sample buffer and analysed by SDS–PAGE and Coomassie blue staining.

### *in vitro* processing assays

The His_6_-MBP-NEDD8-Ub was expressed using vector pLous3 and purified using Ni-NTA-agarose37. 600ng of His_6_-MBP-NEDD8-Ub substrate was incubated with recombinant human NEDP1 protein at 37°C for 30min in 15μl of reaction buffer containing 50mM Tris–HCl (pH 8.0), 50mM NaCl and 5mM b-mercaptoethanol. The reaction was stopped by the addition of 3μl of 6X SDS sample buffer and analysed by SDS–PAGE and Coomassie blue staining^37^. To test the NEDP1 processing inhibitory function of the nanobodies, 600ng of His_6_-MBP-NEDD8-Ub was incubated with 300ng of NEDP1 and increasing amount of the Nbs.

### Immunoprecipitations

U2OS cells either with empty pcDNA3 vector or stably expressing Nb2/Nb9-Flag constructs were lysed in 50mM Tris-HCl, pH 7.4, 100mM KCl, 1% NP-40, 2mM EDTA, protease inhibitors (complete EDTA-free, Roche) for 15min on ice. After centrifugation at 13000rpm for 15min at 4°C extracts were used for immunoprecipitations overnight at 4°C with 20μl of pre-washed Flag beads. Next day beads were washed 3x with 500μl of lysis buffer, resuspended in 50μl of 2xSDS Laemmli buffer and boiled for 5min. Eluates were analysed by western blotting.

### Western blot analysis

For all inputs we routinely used approximately 20μg of total protein. Proteins were resolved in 4–12% precast Bis-Tris gels and transferred onto PVDF membrane using the Bio-Rad Mini Trans-Blot apparatus. Membranes were blocked in 5% milk solution (PBS with 0.1% Tween-20 and 5% skimmed milk) for 1h at room temperature with gentle agitation. Membranes were incubated with the primary antibodies diluted in PBS, 0.1% Tween-20, 3 %BSA and 0.1% NaN3, overnight at 4°C. Membranes were washed 3×10min with PBS, 0.1% Tween-20 and then incubated with the appropriate secondary antibody (1:2000) (Sigma Aldrich) for 1h at room temperature (5% milk). After incubation, membranes were washed 2×15min with PBS, 0.1% Tween-20 followed by 2×5min with PBS. Detection was performed with ECL western Blotting Detection Reagents and membranes were exposed to X-ray Medical Film before being developed.

## Results

### Selection of NEDP1 Nbs

An immune response against recombinant NEDP1 was elicited in llama. The peripheral blood lymphocytes of the llama were used to clone the VHH repertoire in a phage display vector. A VHH library of about 3 × 10^8^ independent transformants was obtained. A colony PCR on 20 clones from the library indicated that 93% of transformants harbored the vector with the right insert size for a VHH gene. Representative aliquots of the library were rescued with M13K07 helper phages, and the highest enrichment was obtained after the third round of panning (100-fold). Finally, based on sequence data, 21 different Nbs (Fig.1A) were identified. The specificity of the isolated Nbs against human NEDP1 and ULP-3 was also determined by ELISA as described in Materials and Methods (Fig. 1A). Further ELISA testing with increasing amounts of coated recombinant NEDP1 confirmed the interaction of a series of isolated Nbs with NEDP1 (Fig. 1B).

**Figure 1.**
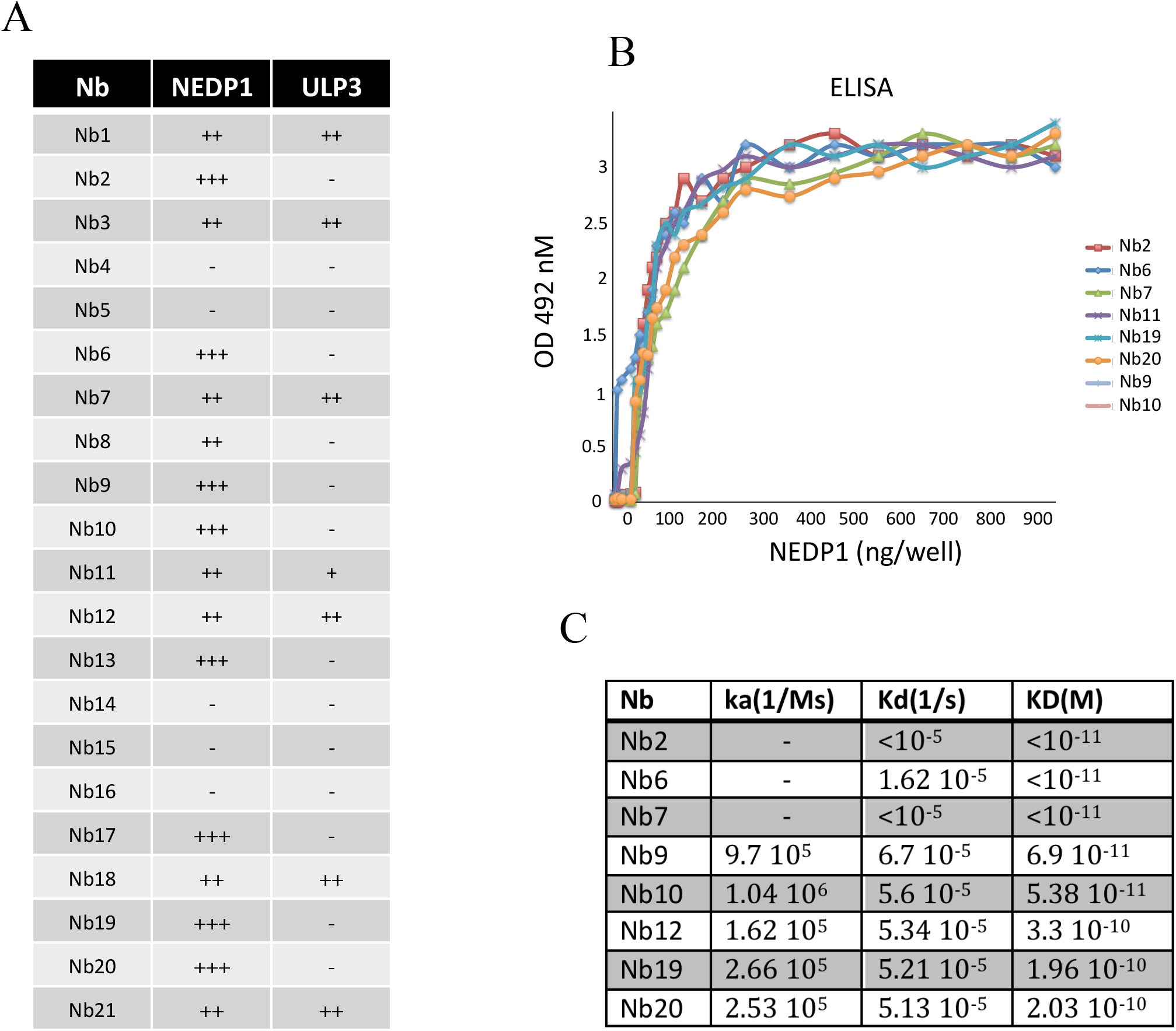
A. Table showing the different Nbs isolated and their binding specificity towards the immunogen human NEDP1 and ULP-3, the *C. elegans* homologue of NEDP1 (− poor/no binding, + indicates binding affinity/specificity). B. ELISA test, performed as described in Materials and Methods, showing the binding of indicated Nbs (300ng) against increasing amounts of coated recombinant human NEDP1. C. Table showing the SPR analysis and binding affinities for indicated Nbs against human NEDP1.

The affinities of selected Nbs against NEDP1 were determined by surface plasmon resonance as described in Materials and Methods. The Nbs were immobilized on CM5 sensor chips and dilutions of NEDP1 were in the mobile phase to record the association kinetics. The binding kinetics between the Nbs and NEDP1 yielded KDs values that vary from 10^−10^M to <10^−11^M (Fig. 1C). Nbs with high affinities, purification yields and specificity for human NEDP1 (Nb2, Nb9 and Nb10) were selected for further functional validation.

### Anti-NEDP1 Nbs inhibit both the processing and deconjugating activity of NEDP1 *in vitro*

NEDP1 was originally discovered as a specific NEDD8 processing enzyme exposing the diGly motif at the C-terminus of NEDD8 but it also promotes deconjugation of NEDD8 from substrates1,2. We first determined the effect of the Nbs on the processing activity of NEDP1 *in vitro*. The assay is based on monitoring the activity of NEDP1 to cleave the His_6_-MBP-NEDD8-ubiquitin fusion. The release of ubiquitin is indicative of the processing activity of NEDP1 (Fig. 2A). As expected wild type NEDP1 can process *in vitro* the His_6_-MBP-NEDD8-ubiquitin fusion (Fig. 2B). Addition of increasing amounts of purified Nb9, completely inhibited NEDP1 processing activity at equimolar amounts (Fig. 2B). Similar results were obtained with Nbs 2 and 10 (Fig. 2C). In contrast, Nb7 which also binds to NEDP1 with high affinity (Fig. 1C), had no effect on the NEDP1 processing activity, strongly indicating that the inhibitory effect of Nbs 2, 9, 10 is specific and not due to a co-purified bacterial contaminating activity.

**Figure 2.**
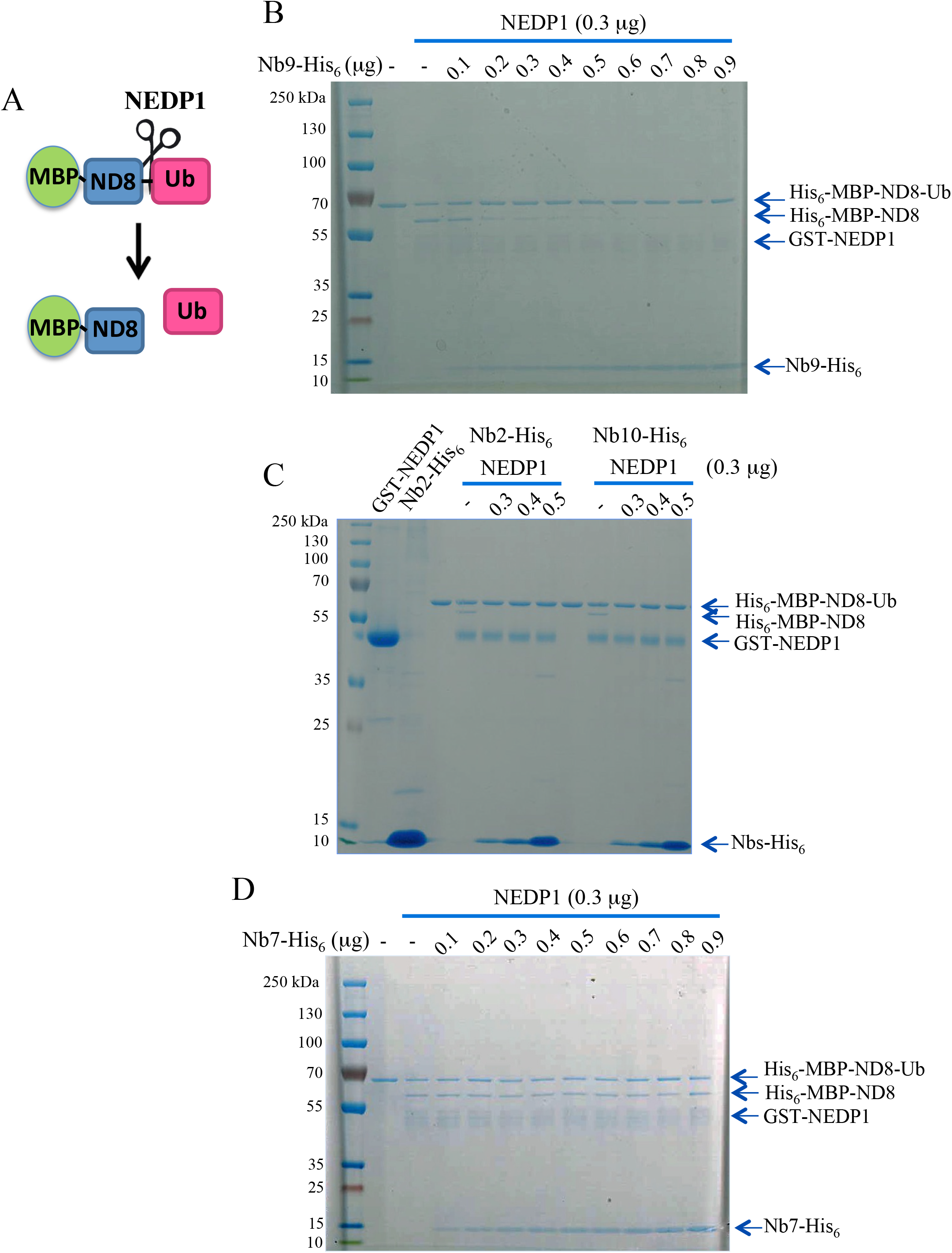
A. Schematic representation of the assay used to determine the effect of Nbs on the NEDP1 processing activity for NEDD8. The His_6_-MBP-NEDD8-ubiquitin fusion protein is incubated with recombinant NEDP1. The production of His_6_-MBP-NEDD8 is indicative of the processing activity of NEDP1. B. The processing assay was performed as described in Materials and Methods were NEDP1 was incubated with His_6_-MBP-NEDD8-ubiquitin alone or in the presence of increasing amounts of purified Nb9. Reactions were analysed by SDS-PAGE and gels were Coomassie blue stained. Arrows indicate the presence of the processing product His_6_-MBP-NEDD8 and of the additional reaction components. C. Assay performed as in B but Nbs 2 and 10 were used instead. In the first two lanes 1μg of NEDP1 or Nb2-His_6_ respectively were loaded as control. D. Similar as in B but Nb7 was used instead.

We next evaluated the effect of the developed Nbs on the deconjugating activity of NEDP1 (Fig. 3A). For this, we used extracts from NEDP1 knockout U2OS cells (H6) that have high levels of mono- and poly-NEDDylated conjugates^19^. Addition of wild type but not catalytically inactive recombinant NEDP1 (Fig. 3B, 3C) dramatically reduces NEDDylation in the extracts indicative of the deconjugating activity of NEDP1. Similarly to the processing assay, addition of purified Nbs 2, 9, 10 at equimolar amounts to the recombinant NEDP1, caused a complete inhibition in the deconjugating activity of NEDP1 (Fig. 3C).

**Figure 3.**
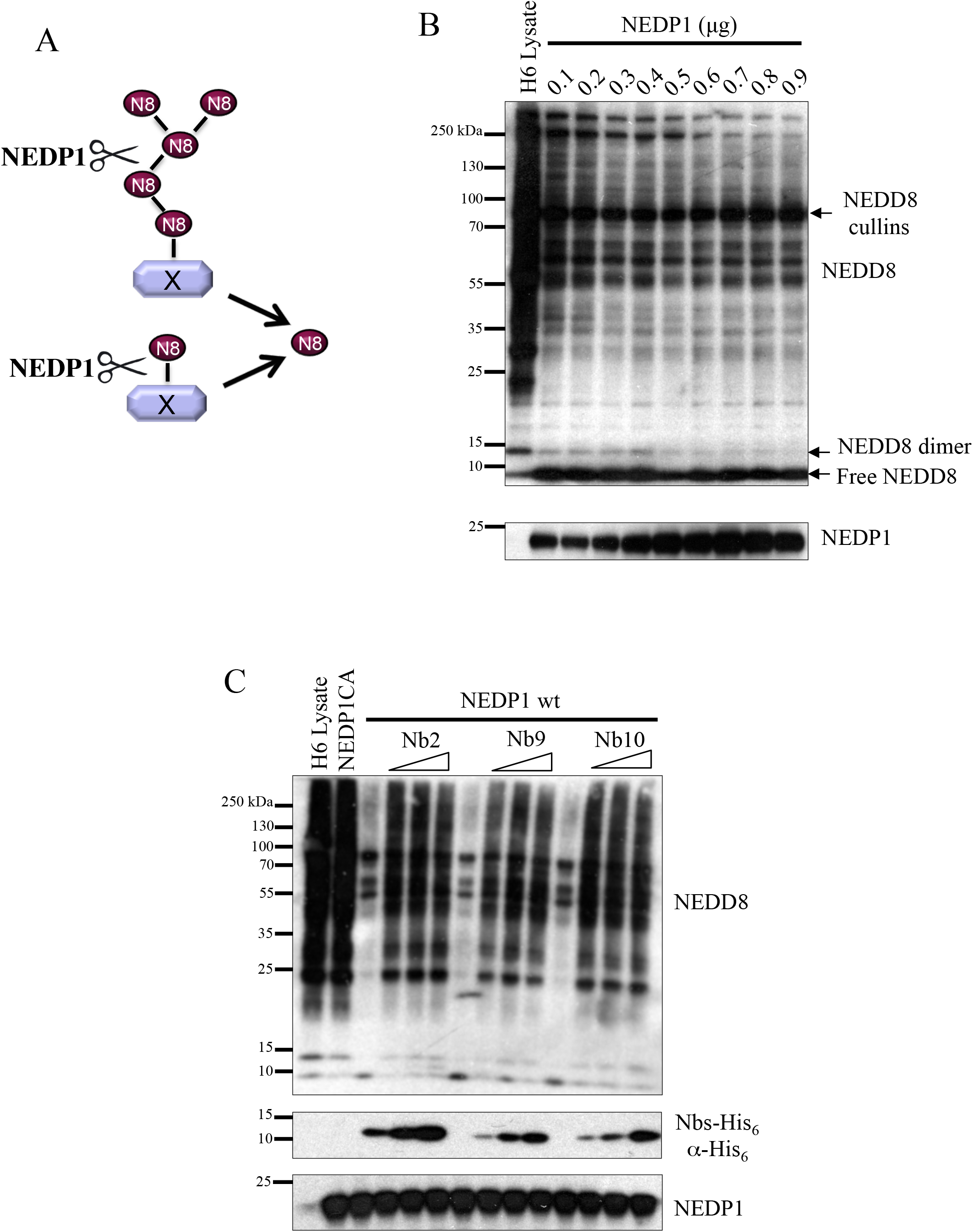
A. Schematic representation of the assay used to determine the effect of Nbs on the deconjugation activity of NEDP1. Extracts from NEDP1 knockout U2OS cells (H6) with high levels of mono- and poly-NEDDylated conjugates were incubated with recombinant NEDP1 in the absence or presence of Nbs. The decrease in global NEDDylation and the accumulation of free-NEDD8 are indicative of the NEDD8 deconjugation from substrates. B. Different amounts of recombinant NEDP1 were added in H6 extracts (20μg total protein) and assay was performed as described in Materials and Methods. Reactions were analysed by western blotting with the indicated antibodies. C. Similar experiment as in B, where 300ng of NEDP1 were added to H6 extracts either alone or with increasing amounts of purified Nb2, 9, 10 (0.1, 0.2, 0.3μg). The catalytic inactive C163A NEDP1 mutant was also used as control for the deconjugating activity of NEDP1.

### Anti-NEDP1 Nbs inhibit NEDP1 activity *in vivo*

To assess the effect of the developed Nbs on the activity of NEDP1 *in vivo* we developed stable U2OS osteosarcoma cell lines expressing Flag-tag versions of Nbs. By anti-Flag immunoprecipitations we found that the expressed Nbs interact with endogenous NEDP1 (Fig. 4A). Consistent with an inhibitory role of the Nbs on NEDP1 function, the levels of NEDDylation are dramatically increased in cells expressing the Nbs, compared to control cells (empty pcDNA3 vector) (Fig. 4A). No effect on the levels of NEDP1 was observed, indicating that the Nbs affect the catalytic activity but not the protein levels of NEDP1 (Fig. 4A). Importantly, both the profile and levels of NEDDylation in cells expressing the Nbs is very similar to those observed in NEDP1 knockout cells^19^, especially on the depletion of free NEDD8, strongly suggesting that the Nbs are potent inhibitors of NEDP1 in cells. No effect on ubiquitination or SUMO-2 conjugation was observed, indicating the specificity of the Nbs towards NEDP1 and the NEDD8 pathway (Fig. 4B). The combination of the above experiments show that the developed Nbs are potent inhibitors of both the processing and deconjugating activity both *in vitro* and *in vivo*.

**Figure 4.**
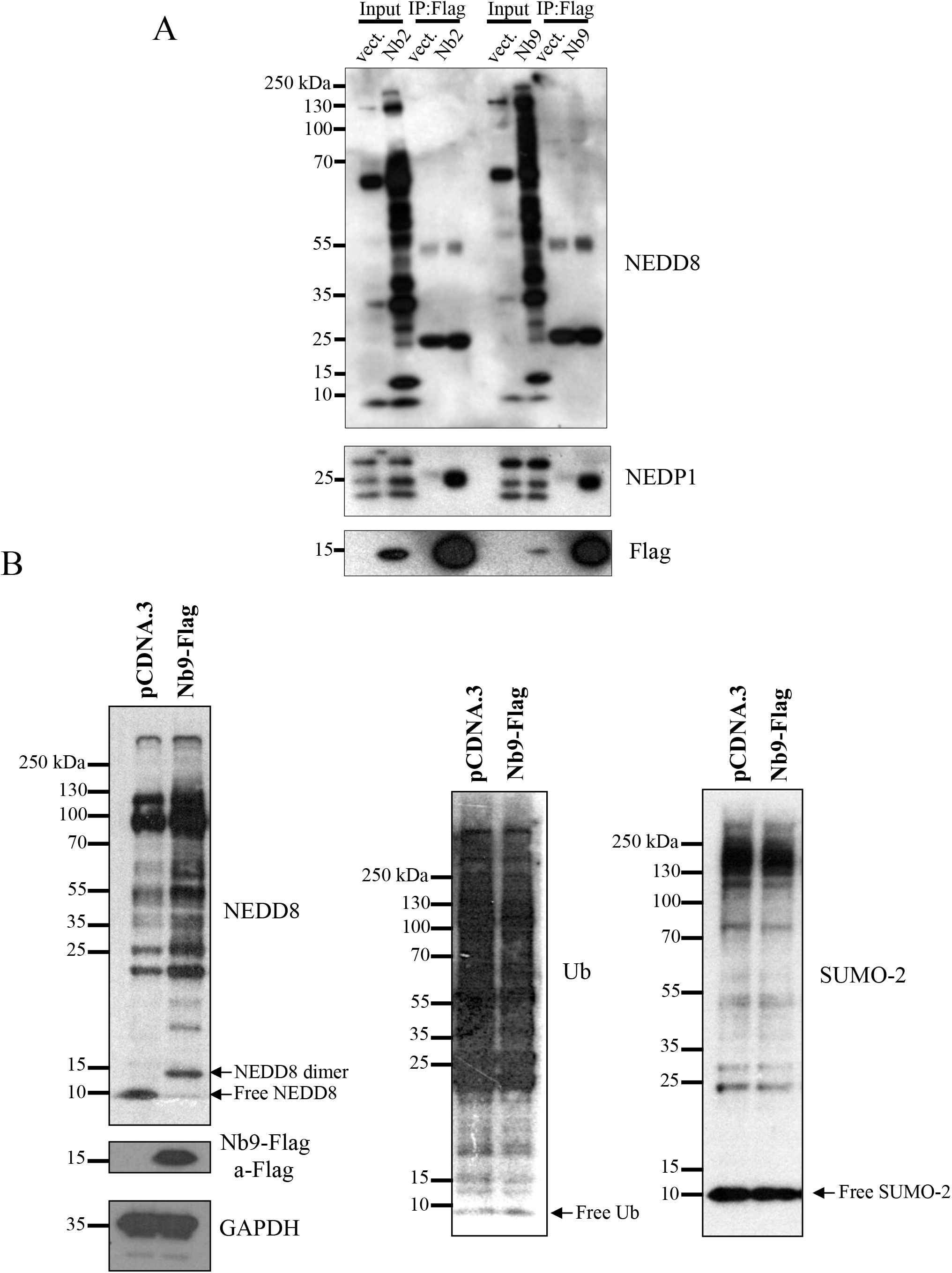
A. Extracts from empty pcDNA3 vector (vect.) or stably expressing Nb2/Nb9-Flag U2OS cells were used for anti-Flag immunoprecipitations. Total cell extracts (input) and eluates were used for western blot analysis with the indicated antibodies. B. U2OS cells described above were lysed in 2xSDS buffer and extracts were analysed by western blotting with the indicated antibodies.

## Discussion

Protein NEDDylation is an important post-translational modification within the family of ubiquitin-like molecules. Components of the NEDD8 pathway are attractive therapeutic targets. Indeed, inhibitors of the NEDD8 E1 activating enzyme are in Phase III clinical trials^1,22^. The deNEDDylating enzyme NEDP1 is now established as a critical regulator of protein NEDDylation, implicated in the control of several cellular processes. Decrease in NEDP1 levels was recently proposed as a molecular basis for the observed increase in protein NEDDylation observed in hepatocellular carcinoma^19,21^. Therefore, the development of inhibitors for NEDP1 will provide important tools for studying the functions for this enzyme. In this study, we describe the development of Nbs that specifically and potently inhibit both the processing and deconjugating activity for NEDP1. The biochemical analysis shows that the Nbs bind to NEDP1 with high affinity at low nanomolar level. While structural studies will provide the molecular basis for this inhibition, we found that the Nbs compete with NEDP1 for NEDD8 binding (data not shown), suggesting that they may block the NEDP1 catalytic site. The *in vitro* analysis shows that the Nbs at equimolar amounts completely block NEDP1 activity. *in vivo*, expression of the Nbs in U2OS cells results in a dramatic increase in protein NEDDylation, which is very similar to that observed in NEDP1 knockout U2OS cells^19^. Based on our experience on siRNA knockdown of NEDP1, over 80% decrease in NEDP1 levels is required to achieve a rather modest increase in NEDDylation^19^. Collectively, the *in vitro* and *in vivo* studies shows that the developed Nbs achieve almost complete inhibition of NEDP1.

NEDD8 processing upon synthesis is essential for the activation and conjugation of NEDD8 onto substrates^1,2^. The observation that similarly to NEDP1 deletion, expression of the Nbs result in increase of protein NEDDylation in cells, suggests that the processing but not the deconjugating activity of NEDP1 is redundant. This notion is further supported from the observations that NEDP1 deletion is not essential in plants, *Drosophila, C. elegans* and human cells^9,11,12,19,20^ and that the defect in asexual development in fungi with NEDP1 (DenA) deletion is independent of the processing activity of NEDP1^10^. The reported proteases with dual specificity for NEDD8 and ubiquitin^1,2^ or currently unknown NEDD8 proteases could mediate NEDD8 processing in the absence/inhibition of NEDP1.

In combination with the available inhibitors of the deNEDDylating activity of the COP9 signalosome subunit CNS5^24^ the developed anti-NEDP1 Nbs provide powerful tools to study the function of these specific deNEDDylating activities. Based on their high potency against NEDP1 and the simplicity to produce them as recombinant proteins, anti-NEDP1 Nbs can be used routinely in cell extracts and native lysis conditions to prevent rapid deNEDDylation due to endogenous NEDP1 activity. Such tools could help the identification and characterization of NEDD8 substrates under entirely endogenous conditions. To our knowledge the described Nbs provide the first-in-class inhibitors for NEDP1.

## Notes

#### Summary of Updates

Protein NEDDylation emerges as an important post-translational modification and an attractive target for therapeutic intervention. Modification of NEDD8 onto substrates is finely balanced by the co-ordinated activity of conjugating and deconjugating enzymes. The NEDP1/DEN1/SENP8 protease is a NEDD8 specific processing and deconjugating enzyme that regulates the NEDDylation mainly of non-cullin substrates. Here, we report the development and characterisation of nanobodies as first-in-class inhibitors for NEDP1. The nanobodies display high-affinity (low nM) against NEDP1 and specifically inhibit NEDP1 processing activity in vitro and NEDP1 deconjugating activity in tissue-culture cells and in cell extracts. We also isolated nanobodies that bind to NEDP1 with high-affinity but do not affect NEDP1 activity. The developed nanobodies provide new tools to study the function of NEDP1 and to prevent deNEDDylation in cell extracts used in biochemical assays.

